# Ancestry informative alleles captured with reduced representation library sequencing in *Theobroma cacao*

**DOI:** 10.1101/407387

**Authors:** Jaime A. Osorio-Guarín, Corey R. Quackenbush, Omar E. Cornejo

## Abstract

As the source of chocolate, cacao has become one of the most important crops in the world. The identification of molecular markers to understand the demographic history, genetic diversity and population structure plays a pivotal role in cacao breeding programs. Here, we report the use of a modified genotyping-by-sequencing (GBS) approach for large-scale single nucleotide polymorphism (SNP) discovery and allele ancestry mapping. We identified 12,357 bi-allelic SNPs after filtering, of which, 7,009 variants were ancestry informative. The GBS approach proved to be rapid, cost-effective, and highly informative for ancestry assignment in this species.

## Introduction

*Theobroma cacao* L, the tree from which chocolate is confectioned, originated in the Amazonian basin [1] and has become a globally cultivated crop in recent times [2]. There are more than 20 species within the *Theobroma* genus, which belongs to the Malvaceae family [3]. *T. cacao* is the only species that is cultivated extensively because it can be manufactured into chocolate, liquors, confections, cosmetics, and animal feed [4,5]. Chemical and archaeological evidence shows that human settlements have been consuming cacao since 1,400 BC [6,7]. Since 900 BC *T. cacao* has been widely cultivated in Mesoamerica, which is considered the center of domestication. Recent genetic work has supported these findings suggesting that domestication of cacao occurred 3.6 Kya [8]. The domestication of cacao has resulted in the development of an important crop that sustains the livelihood of communities throughout Central and South America, Asia, and Africa [1,9,10], and is the principal income for about six million smallholder farmers globally that produce up to 90% of the world’s cocoa [11].

Cacao is a diploid species with a chromosome complement of 2n=20 [12]. Recently, genomic resources have been developed to facilitate the study of the species evolutionary history, the inference of the genetic basis of phenotypes of interest, and the development of marker assisted breeding programs. There are two reference genomes that have been sequences from different cultivars, the B97-61/B2 genome (Criollo), with an estimated size of 409 million base pairs (Mbp) [13] and the Matina 1-6 genome (Amelonado), with an estimated genome size of 445 Mbp [14]. In addition, Cornejo et al. 2017 [8], performed whole genome sequencing on 200 cultivars in order to understand the domestication process in this species. Recently, approaches to infer genetic diversity in *T. cacao* have been completed using a small panel of single nucleotide polymorphisms (SNPs, ranging from 87 to 6,000) developed with a Fluidigm array system [15–17]. Also, using the information generated from a subset of the 200 cultivars project, an Ilumina Infinity II genotyping array of 13,530 markers was recently developed [18]. However, it is well known that genotyping strategies based on arrays are biased towards the capture of known specific loci in reference populations which introduces ascertainment bias in the genotyping of samples from different origins [19]. There is a growing interest in developing strategies to rapidly genotype accessions at a reduced cost in order to perform extensive association studies and better identify accessions with specific markers that will accelerate assisted breeding programs.

The generation of genomic resources in cacao has benefited from advances in sequencing chemistry, cost reductions, and increased computational capacity to analyze data [20]. Yet, despite the general reduction of costs, the sequencing of complete cacao genomes remains a costly approach for breeders and researchers. Due to the high cost of whole genome sequencing, genotyping arrays are typically used. However, genotyping arrays suffer from the aforementioned issues of ascertainment bias, which hinders the identification of novel untapped genetic variation in the species. It has been shown that genotyping-by-sequencing (GBS) is a cost effective and efficient methodology for the generation of a large number of SNPs, while avoiding ascertainment biases [21,22]. The genetic information generated by GBS has been shown to facilitate robust studies of genetic diversity, population structure, phylogeny, and association in a large number of plant and animal species [23–29] but have never been applied to the study of *T. cacao*. The original protocol proposed for the digestion and generation of reduced representation genomic libraries used the *ApeKI* enzyme [21]. However, Cooke et al. (2016) [30], demonstrated that the use of enzymes that cut far from the recognition site, such as *BsaXI*, increases the diversity of the library and therefore the number and quality of reads sequenced. In the original protocol the restriction enzymes would produce libraries with nearly identical initial sequence causing saturation of the image capture during sequencing. To avoid this issue, the original protocol incorporated the barcode in one of the adapter sequences upstream of the cut-site to introduce variation and minimize the problems associated with saturation [31]. In the proposed protocol (taken from Cooke et al. 2016 [30]), the chosen enzyme generates libraries in which the initial sequence is random due to the fact that the enzyme cuts 14 base pairs (bp) away from its recognition site.

In this work, we adapt and modify an existing GBS method [30] to generate a large number of ancestry informative SNPs in cacao. We further show that the data generated produced genetic diversity estimates consistent with what has been shown in previous work. We also provide an estimate of the proportion of novel versus common SNPs (based on the relative frequency of SNPs) that can be generated using the proposed methodology. Due to the lower cost of generating this type of data we propose that this strategy could be widely used to perform genetic diversity analysis on thousands of samples, facilitating association mapping studies aimed at identifying the genetic basis of important agronomic traits in cacao such as disease resistance and bean quality.

## Methods

### Plant materials

A total of 30 accessions from the germplasm bank from Cocoa Research Centre at the University of West Indies in Trinidad **(S1 Table)** were included in the analysis. Young leaves from one individual per accession were sampled in Trinidad and sent to the Cornejo Lab at Washington State University. Small strips (100 mg) were cut with sterile scissors (washed with 1% hydrochloric acid and ethanol prior to and in between sample processing), placed in 1.5 ml Eppendorf tubes and stored at −80 °C for DNA extraction. Five samples (GU 175/P, ICS 1, IMC 67, M 8[SUR], SCA 6) from the 200 genomes project were genotyped in this study in order to compare the concordance between SNPs identified using GBS to those identified using full genome sequencing [8].

### DNA extraction and GBS library preparation

Total DNA was isolated from 100 mg of young leaves collected from each accession using a DNeasy Plant Mini Kit (QIAGEN, Germany) according to manufacturer’s instructions. The final elution volume was adjusted to 60 μl with 1X TE solution buffer. Integrity of the total DNA was checked on a 0.7% agarose gel. DNA concentration was measured using a Qubit Fluorometer (Life Technologies, Thermo Fisher Scientific Inc., Waltham, MA, USA) with a dsDNA HS Assay Kit (Life Technologies). All samples were normalized to the same volume (50 μl) containing 500 ng of DNA. For each sample, genomic DNA was digested with 2.4 units of *BsaXI* (New England Biolabs, NEB, R0609S) at 37 °C overnight in a 100 μl reaction buffered with Cut Smart NEBuffer (NEB, B7204S). After digestion, we performed a 0.7X size selection using AMPure XP beads to remove any large undigested fragments. The DNA was then prepared for sequencing using an NEBNext Ultra DNA Library Prep Kit for Illumina (NEB E7370S; version 1.2) with the following modifications. Adapter ligation was followed with a bead clean up without size selection. Library enrichment was accomplished using 10 cycles of PCR amplification. Finally, we evaluated the distribution of fragment lengths for each library using an Agilent 2100 BioAnalyzer. Samples were sequenced on an Illumina HiSeq 2500 (Illumina Inc. San Diego, CA), with paired-end 100 bp reads.

### SNP discovery and data processing

Raw sequence reads were demultiplexed and converted to Fastq files by the Spokane Sequencing Core at Washington State University using Casava v1.8.2 (Illumina Inc.). SNP quality control and trimming were conducted using FastQC [32] and Trim Galore v0.5.0 [33], respectively. Reads were trimmed to remove 3’ bases with a quality score less than 25. Reads in which complete or partial adapter sequences were identified were trimmed accordingly. Reads less than 60 bp in length after trimming were removed. The trimmed sequences were aligned to the Matina reference genome using BWA v0.17.0 [34] with the default parameters (except for mismatch penalty -B 6). We performed quality assessment of the BAM files, validation read groups, removal of PCR duplicates, and correction of mapping quality assignment with Picard v2.18.9 [35]. The genome analysis toolkit (GATK) v3.8.1 [36] was used to perform local realignment and base quality recalibration. We performed SNP calling using the unified genotyper module in GATK to facilitate comparison with previous work. Hard filters were applied to the data (quality by depth higher than 2, strand bias FS 60, Root mean square of mapping quality 40.0) and only SNPs that passed the filters were retained.

For population structure analysis we used VCFtools v0.1.13 [37] to performed additional variant filtering in which only bi-allelic variants with a minimum minor allele frequency (MAF) greater than 0.05 were retained. Coverage statistics were obtained by analyzing these loci with Plink-Seq v0.10 [38]. To analyze the types of mutations we used the program SnpEff with the genome built in database [39]. The R scripts are provided in the GitHub of the project.

### *In silico* digest

*In silico* digestion of the cacao chromosomes were performed with the *T. cacao* cv. Matina reference genome [14]. To identify all restriction cut site positions for *BsaXI*, the program RESTRICT from the Emboss package v6.5.7.0 was used [40]. Digested fragments were ordered per chromosome and then were summed using two size ranges (200-700 bp and 200-1000 bp). The desired range for the libraries to be sequenced on an Illumina HiSeq is between 200-700 bp, however fragments of up to 1000 bp can be sequenced [41]. We report the distribution of the fragments as a histogram and the location of the *in silico* digested fragments along the genome.

To estimate the number of fragments produced from the *BsaXI* GBS experiment and to compare them with the *in silico* predictions, the alignment files were analyzed using INTERSECT included in BEDtools v2.27.0 [42]. Using the output files from this command we estimated the number of *BsaXI* fragments for all chromosomes and compared the results with those from the *in silico* digestion.

### Genetic diversity

Observed heterozygosity (Ho), expected heterozygosity (He), and polymorphism information content (PIC) were calculated with Power Marker v1.25 [43]. Tajima’s D was computed with VCFtools v0.1.13. We also estimated allele frequencies using VCFtools for each of the sites that met the following criteria: i) bi-allelic and ii) no missing data. For each site we used folded site frequency spectrum (SFS) for the cacao GBS dataset, which describes the genetic variation as a distribution of allele frequencies across the genome. The site frequency spectrum was generated in R.

### Admixture and population genetic structure analysis

The population analysis was carried out on the filtered set of SNPs. We analyzed the ancestry and admixture patterns for the 30 unrelated accessions using cacao samples recently sequenced [8]. The intersected dataset of MAF filtered SNPs between our cacao GBS dataset (test set) and the 79 individuals from the whole sequence dataset (reference set) consisted of 5,225 SNPs. We ran ADMIXTURE v1.3 [44] in supervised mode assuming ten populations, following Motamayor et al. 2008 [45] and Cornejo et al [8]. Principal component analysis (PCA) was performed with the algorithm implemented in EIGENSOFT [46]. All figures were generated using R version v3.5.1 [47]. The precision of the ancestry analysis was evaluated by obtaining bias and standard error for each ancestry out of a bootstrap with 2000 pseudo-replicates in ADMIXTURE v1.3 [44].

## Results

### *In silico* restriction enzyme digestion

To determine i) what the expected number of fragments to be sequenced was; and ii) to infer how many of the sequenced fragments matched locations in the genome that were expected from the digestion, we evaluated the distribution of fragments throughout the genome (**S1 Fig**). Those sequenced regions that did not match the locations identified from the *in silico* digest were considered off target. The *in silico* digestion was done considering two sets: 1) lengths ranging from 200 to 700 bp generated a total of 5,479,477 bp **(Fig 1A**) and 2) lengths ranging from 200 to 1000 bp generated a total of 10,217,990 bp **(Fig 1B)**. These represent 1 and 3% of the genome, which suggests that *BsaXI* is an appropriate cutter. We used the larger range to evaluate overlap of sequenced target and predicted target.

**Fig 1.** *In silico* analysis of restriction enzyme sites in the *T. cacao* genome. /bold> (**A)** Number of fragments computed in the size range between 200 and 700 bp. **(B)** Number of fragments computed in the size range between 200 and 1000 bp.

The comparison between the predicted fragment-size distribution with the fragments produced through our assay **(Fig 2)** suggests high variability in the accuracy of sequenced targets across samples. Despite the high variability, 73% of the samples had an overlap of over 57% with the *in silico* digest, and 53% of samples had an overlap over 76% with the expectation

**Fig 2.** Assessment of sequencing on target per sample. The orange bar corresponds to the total number of fragments that are predicted to be digested *in silico* within the length distribution of 200 to 700 bp. Blue bars correspond to the number of sequenced fragments that meet our callability criteria overlapping with those predicted *in silico*.

### SNP Discovery

We obtained a total of 366 million paired-end 100 bp reads. The number of raw sequence reads per individual ranged from 0.4 million to 52.2 million. After removal of low-quality sequences and adapter trimming, 341 million reads remained. Of these reads, 322 million aligned to the Matina reference genome, and were used for SNP calling. The mean read depth ranged from 8.9 to 160.1. The observed differences in read depth across samples is most likely due to differences in quality and/or variations in barcode efficiencies. Individual data information is shown in **Table 1**.

**Table 1.** Summary statistics of sequenced data per individual.

| Accession Number | Reads raw-data (Millions) | Reads after trimming (Millions) | Mapping reads (and percent) | Depth |
| --- | --- | --- | --- | --- |
| CL 10/11 | 0.53 | 0.50 | 0.49 (97.6%) | 15.2 |
| CL 10/3 | 4.8 | 4.7 | 4.5 (96.8%) | 67.1 |
| CLM 100 | 14.9 | 14.6 | 14.2 (97.4%) | 131.2 |
| GS 6 | 14.5 | 14.1 | 13.8 (97.8%) | 122.9 |
| GS 61 | 10.2 | 9.9 | 9.5 (96.4%) | 104.2 |
| GU 175/P_1 | 5.3 | 5.1 | 5.0 (96.7%) | 73.6 |
| GU 175/P_2 | 1.1 | 1.04 | 1.01 (96.6%) | 9.0 |
| ICS 1_1 | 3.2 | 3.1 | 3.0 (96.9%) | 52.6 |
| ICS 1_2 | 4.9 | 4.7 | 4.5 (95.6%) | 86.7 |
| ICS 28 | 1.9 | 1.88 | 1.82 (96.8%) | 26.6 |
| ICS 87 | 3.1 | 1.67 | 1.62 (96.9%) | 21.6 |
| IMC 6 | 7.9 | 5.3 | 5.1 (95.7%) | 71.6 |
| IMC 67_1 | 1.91 | 1.90 | 1.83 (96.4%) | 26.6 |
| IMC 67_2 | 0.4 | 0.36 | 0.35 (97.6%) | 8.9 |
| LCT EEN23 | 10.4 | 8.3 | 8.2 (98.2%) | 126.6 |
| LCT EEN246 | 8.2 | 7.8 | 7.6 (97.1%) | 100.4 |
| LCT EEN31 | 7.8 | 6.2 | 6.0 (96.3%) | 74.8 |
| LP 3/29[POU] | 7.4 | 5.2 | 5.0 (96 %) | 62.3 |
| LZ 33 | 8,0 | 6.2 | 6.0 (97.4%) | 66.3 |
| M8 [SUR]_1 | 4.6 | 3.4 | 3.3 (96.6%) | 52.1 |
| M8 [SUR]_2 | 15.2 | 14.8 | 14.4 (97.4%) | 125.9 |
| NA 189 | 7.2 | 5.2 | 5.1 (96.9%) | 72.4 |
| NA 26 | 12,0 | 11.7 | 11.2 (95.9%) | 128.5 |
| NA 68 | 6.6 | 6.4 | 6.2 (96.7%) | 86.6 |
| RIM 113[MEX] | 10.8 | 10.5 | 10.1 (96.1%) | 39.9 |
| SCA 6_1 | 2.6 | 2.5 | 2.4 (96.4%) | 39.9 |
| SCA 6_2 | 8.4 | 8.1 | 7.9 (96.6%) | 101.2 |
| SLC 18 | 13.9 | 13.1 | 12.7 (97.1%) | 123.9 |
| SLC 8 | 20.6 | 19.5 | 18.8 (96.6%) | 141.9 |
| UF 38 | 52.2 | 50.9 | 49.05 (96.2%) | 160.1 |

**Table 2.** Summary of polymorphisms identified in cacao dataset.

| Total number of polymorphisms |  |  |
| --- | --- | --- |
| **SNPs** | 12,357 | 100% |
| **Transition** |  |  |
| **A/G** | 3,856 | 31.2% |
| **C/T** | 3,894 | 31.5% |
| **Transversion** |  |  |
| **A/C** | 1,243 | 10.1% |
| **A/T** | 1,403 | 11.3% |
| **C/G** | 758 | 6.2% |
| **G/T** | 1,203 | 9.7% |
| **Ti/Tv ratio** | 1.79 |  |

A total of 12,357 SNPs were identified in the collection. The SNPs were classified into transitions (Ti) and transversions (Tv) based on nucleotide substitution. The number of A/G transitions was very similar to the number of C/T transitions. The numbers of A/C and A/T transversions were relatively higher than the C/G and similar to G/T transversions. Transitions are the most common type of nucleotide substitutions [48]. In this dataset 62.7% of the base changes were transitions and 37.3% were transversions

### Genetic diversity analysis

The dataset was reduced to 7,009 SNPs after excluding markers showing: 1) more than 20% missing data, 2) a MAF ≤ 0.05, 3) multi-allelic sites, 4) insertions and deletions. Expected heterozygosity (He) (mean 0.272), Ho (mean 0.200) and PIC (mean 0.226) values estimated per position are listed in **S2 Table**. He, Ho and PIC ranged from 0.117 to 0.500, from 0 to 0.437, and from 0.374 to 0.375, respectively. Tajima’s D was 0.458, indicating an excess of intermediate frequency alleles that could result from demographic processes such as population bottlenecks, population subdivision or migration. The SFS showed a significantly high proportion of rare alleles (45.3% of identified alleles). These results quantitatively show the proportion of rare variants that would be missed if genotyping arrays were employed for detecting and characterizing genetic variation **(Fig 3)**. The SFS shows a variable pattern of frequencies of alleles at intermediate minor allele counts, which is consistent with unaccounted for population structure.

**Fig 3.** The site frequency spectrum of *T. cacao* samples used in the study. The orange bars correspond to alleles found in one, two or three chromosomes in the sample, considered to be low frequency in this study. The green bars correspond to alleles with a minor allele count of 4 or more and are considered common variants for the purpose of this experiment. Conservative estimates suggest that nearly 50% of the SNPs are rare and would be missed if an array were used to genotype these accessions.

### Admixture and population genetic structure analysis

The filtered set of SNPs was used for admixture and population structure analysis. The 30 accessions were compared with ten reference populations that are related to the cacao groups previously defined [45]. Fifty percent of analyzed individuals were assigned to unique ancestry in the Curaray, Nanay, Contamana, Amelonado, Nacional, Guianna and Iquitos populations. The other half of the individuals presented a variety of mixed ancestries **(Fig 4)**. The analysis of bias and error supports a high level of precision in the assignment of ancestry for the majority of the samples (**S2 Fig A**). For the most part, bias in the assignment was lower than 5%. However, LCTEEN23 presented a relatively higher bias (10%) in the assignment to Nacional ancestry (**S2 Fig B**). Standard error (SE) is relatively low overall with the majority being below 0.05. Sample LCTEEN23 presented a relatively higher SE in the assignment of Nacional ancestry (**S2 Fig C**).

**Fig 4.** Supervised ADMIXTURE using ten reference populations. Each bar corresponds to an individual and the height to the proportion of ancestry explained by one of the 10 previously described populations. The first uniformly colored 79 columns correspond to individuals in the reference set. The remaining 30 columns show the proportions of ancestry assignment for each one of the newly sequenced cacao samples with our GBS approach.

In addition, pairwise allele (SNP) sharing distances were computed between all samples and reference populations and PCA was used to illustrate the resulting pairwise distances **(Fig 5)**. Plotting the two first axes showed that they capture the previously described genetic groups with distinct clusters reflecting the different origin in cacao today. The first principal component (13.74% of the variation) showed a clear relationship between the samples and the Criollo and Amelonado ancestral populations whereas the remaining populations (Curaray, Nanay, Contamana, Maranon, Nacional, Gianna, Iquitos, and Purus) separated along the second principal component (11.77% of the variation).

**Fig 5.** Principal component analysis of reference and cacao GBS samples. PCA plots of the same cacao collection according to subgroups, as identified by ADMIXTURE software. Although reference samples are closer together, the newly sequenced individuals are placed close to other individuals showing similar ancestry. Admixed individuals in the new cacao GBS set are more variable and overdispersed when compared to the reference set.

## Discussion

Marker assisted breeding in cacao currently faces a great challenge in the development of alternative fast, cheap, and accurate genotyping strategies for a large number of accessions. The need for inexpensive and accurate genotyping strategies is especially true when considering that most cacao producers are located in developing countries with limited resources. Some of the technologies developed for array genotyping, which have been advanced by us and others, work well for the identification of known variants but are limited in the ability to discover new and potentially informative variants. Genotyping-by-sequencing is a high-throughput, low-cost technology, which is useful for variant identification in diverse species and populations and allows for the discovery of previously unidentified variants [21,49]. In this work, we have adapted and modified a GBS genotyping assay for cacao. The most important changes to the library preparation are: 1) the use of the restriction enzyme *BsaXI* that cut 14 bp away from the recognition motif [30], 2) the use of the Illumina Y-adapters allowing for PCR incorporation of dual-indexed barcodes, which facilitates large-scale, inexpensive multiplexing, and 3) the efficient cleaning of non-ligated adapter excess and PCR duplicates using cleaning beads, which is a widely used technique already employed in previous work [8].

There was variation across samples in the number of mapped reads that overlapped with the fragments predicted from the *in silico* digestion. The performance of our genotyping in on-target regions is similar to other studies [21], while we produced a large number of reads per sample (0.5 M – 14.3 M reads), only a fraction overlapped with predicted regions for a number of samples. The lack of overlap for some samples suggests that a large number of fragments sequenced were off target. For example, GU 175.2 had a low percent of overlap (5%), whereas UF 38 had 95% overlap with the *in silico* digest (**Fig 2**). The number of sequenced fragments overlapping with *in silico* predicted regions is likely explained by variations in DNA quality that result from fragmentation during DNA extraction, prior to the digestion.

We identified a large number of variants, 12,357 SNPs, that were retained after filtering. Estimates of genetic variability are comparable with those estimated in previous studies of cacao [18]. Enhanced SNP discovery, SNP quality, production steps, and optimization of parameters improved the SNP detection when compared to those obtained from 96 SNPs from small Fluidigm arrays or the 6K SNP array previously reported [15–17]; or even from the 15K SNP arrays considering the fact that a large proportion of SNPs cannot be identified with array systems. According to Romay et al. 2013 [50], one important advantage of GBS over the SNP array systems, is the MAF distribution. We are interested in the identification of novel genetic variation that might prove to be useful in the identification of traits of interest and much of this genetic variation may not be captured with genetic arrays. The allele frequency distribution of variants identified using GBS followed an expected pattern with an excess of rare variants **(Fig 3)**. More than 45% of bi-allelic variants were rare and the distribution of intermediate variants revealed signatures of population structure.

The mean expected heterozygosity value was higher than the observed heterozygosity, indicating a deficit of heterozygotes. Compared with recent published genetic diversity studies in cacao [16,17,51], our results present slightly lower values of He, Ho and PIC, probably due to the high presence of homozygous genotypes of the Criollo and Amelonado populations and the impact of a well-known effect of allelic dropout in GBS datasets [30,52].

The pattern of population structure was further validated when ancestry and structure were analyzed together with the reference panel set. To study the ancestry of the newly sequenced samples 5,225 bi-allelic SNPs were successfully intersected with a reference panel, which consisted of 79 fully sequenced genomes. The samples used for the reference set were recently sequenced [8] and represent the 10 different genetic groups from the Upper Amazon, Lower Amazon, Orinoco, and Guyana [45]. Our ancestry characterization of the newly sequenced 30 samples was consistent with the known passport data for the accessions **(S1 Table)**. More specifically, accessions NA 26 and NA 189 were successfully assigned to the Nanay group and both SCA 6 accessions were correctly assigned to Contamana, showing that the procedure allows us to correctly identify their population of provenance. In the case of admixed individuals, we could correctly assign ancestry to individuals with known admixture patterns. For example, for ICS or UF individuals which are known to be a mix between Criollo and Amelonado. These results make us confident that the generated data and procedure could be used to disentangle more complex patterns of genetic admixture like those obtained for previously uncharacterized individuals such as LCT EEN 246. The accession LCT EEN 246 is comprised of Criollo, Marañon, Iquitos, and Contamana ancestry, which could open new possibilities for studying traits of interest.

Overall, the observed patterns of ancestry are consistent with a known history of the hybridization process in cacao breeding programs and natural hybridization [8]. Studies with a limited set of markers have already hinted at the possibility of further substructure within the ten main populations described for cacao [45,53]. Our results show the usefulness of GBS to address these questions.

## Conclusions

We have shown the increased potential for the identification of ancestry informative and novel variants, which are useful for addressing questions about population demographic history and genomic variation in *Theobroma cacao*. We were able to identify 12,357 SNPs markers from only 30 samples, out of which 7,009 could be used for ancestry determination. There is a need to increase our knowledge about cacao accessions throughout the world to produce high yielding and disease resistance genotypes. Our results have practical implications for the development of strategies for genotyping large numbers of accessions with the aim of performing association mapping studies. In our future work, the GBS technique will be refined to include a two digestion enzymes protocol to genotype larger cacao collections and increase the number of common targets/SNPs identified with the method.

## Acknowledgments

We thank Path Umaharan, Lambert Motilal, and Annelle Holder-John for providing the germplasm material and associated information used in this study. The authors thank Joanna Kelley for assistance in revising the final version of the manuscript. The funders had no role in study design, data collection and analysis, decision to publish, or preparation of the manuscript. Jaime Osorio was supported through funds assigned from Agrosavia-Colombia to the Cornejo Lab to work on the library preparation of the samples.

## Data availability statement

Raw sequence reads (data to SRA) is available at NCBI Bioproject ID No. PRJNA486150. The scripts are available at GitHub repository (https://github.com/jaog224/cacao_gbs).

## Supporting information

**S1 Table. Summary of the accessions of the *Theobroma cacao* collection used in this study.**

**S2 Table. Summary statistics of genetic diversity of the 7,009 SNPs.**

**S1 Fig. Distribution of *BsaXI* enzyme cut sites across the *T. cacao* cv. Matina genome.**

**S2 Fig. Precision of the ancestry analysis using 200 bootstrap pseudoreplicates. (A)**

ADMIXTURE bar plot, **(B)** bias bar plot and **(C)** standard error for each ancestry.

